# Image-based Disease-wide Association Study via Self-supervised Learning links Abdominal MRI Features to 158 Diseases

**DOI:** 10.64898/2026.07.23.740409

**Authors:** Seyed Mousavi, Jatin Arora, Richa Agarwal, Anja Fischer, Stefanie Mueller, Zhihao Ding, Johann de Jong, Ke Yuan

## Abstract

Medical images contain rich phenotypic information that is often not fully captured by manual clinical assessment. Here, we present a systematic framework for extracting such information and conducting image-based disease-wide association studies (iDWAS) using self-supervised learning (SSL). We applied this framework to abdominal MRI data from the UK Biobank and evaluated associations between MRI-derived features and 562 diseases. Features were learned using a VICReg-based SSL model and tested for disease associations using logistic regression models. We identified 158 diseases that were significantly associated with the MRI-derived features, including the ones which are not directly linked to abdominal anatomy. Focusing on Metabolic dysfunction–associated steatohepatitis (MASH), we showed that the MRI-derived features captured disease-relevant information and separated MASH cases from controls better than established biomarkers such as PDFF and Iron-cT1. These findings highlight the ability of SSL to uncover clinically meaningful signals from routine imaging data. The proposed framework is broadly applicable to other imaging datasets and modalities, enabling more systematic approaches to incidental and early disease detection. The trained model and analysis pipeline are publicly available at https://github.com/srm2022/iDWAS.

**Background:** Medical images, often analyzed manually by clinicians, contain valuable information that may not be fully captured by the human eye. Machine Learning (ML) has demonstrated significant potential in extracting this information, allowing us to enhance our understanding of underlying pathologies. We present a systematic framework for extracting such information and performing image-based disease-wide association studies (iDWAS) using self-supervised learning. We then applied the framework to abdominal Magnetic Resonance Imaging (MRI) from UK Biobank (UKB) and investigated the associations between MRI-derived features and 562 diseases. Finally, we delved deeper into the observed associations for Metabolic dysfunction–associated steatohepatitis (MASH), a prevalent yet underdiagnosed condition with a high unmet need for non-invasive diagnostic tools.

**Methods:** We extracted features from MRI images using VICReg-based self-supervised learning and modeled the associations of the extracted features with 562 diseases using logistic regression. Using the likelihood ratio (LR) test, we then assessed how much additional variance can be explained by the MRI-derived features relative to the variance explained by a null model including only common confounders such as age, gender, and BMI.

**Results:** Of these 562 diseases, we identified 158 diseases to be significantly associated with the MRI-derived features. While many of them (e.g. MASH) are known to primarily emerge in the abdomen, others (e.g. neuropsychiatric diseases) are not. Taking MASH as a use-case and adjusting for MASH-specific confounding, we then showed that our MRI-derived features captured additional MASH-relevant information compared to standard MRI-derived biomarkers of liver health such as proton density fat fraction (PDFF) and iron-corrected T1 (Iron-cT1). The trained SSL model and analysis pipeline are available at https://github.com/srm2022/iDWAS.

**Conclusions:** Our results illustrate (1) how MRIs from one organ of the body can inform about the diseases primarily rooted in other organs of the body, reflecting patients’ overall health status, and (2) how SSL can extract more information relevant for specific diseases compared to current state-of-the-art biomarkers. Our framework can be applied to other MRI datasets as well as other imaging modalities with minimal adjustment, providing opportunities for the development of more systematic approaches to incidental diagnosing, i.e. (early) detection of diseases falling outside the scope of the original imaging procedure.

## Introduction

Imaging modalities such as Magnetic Resonance Imaging (MRI) and Computed Tomography (CT) scans contain valuable biological information that is often not fully captured by human interpretation. Traditional approaches relying on manual feature engineering can be labor-intensive, susceptible to human biases, and may fail to extract the full richness of the data, thereby limiting their ability to uncover novel patterns or achieve high predictive performance.

To address these limitations, self-supervised learning (SSL) has recently been employed to automatically extract informative representations from medical images in the form of feature vectors, which can then be utilized in downstream tasks such as diagnosis and disease subtyping. For example, SSL-based approaches have been used to predict liver fat from liver ultrasound scans (Mohit et al. 2025), to identify genetic associations of eye diseases from Optical Coherence Tomography (OCT) scans in the UK Biobank (UKB) (Xie et al. 2024), or to study the genetic architecture underlying variety of diseases using brain and abdominal MRI scans (Sankarapandian et al. 2024). However, such studies have so far focused on specific organs or diseases, rather than exploring associations across a spectrum of conditions in an unbiased manner. Shang et al. (2022) investigated associations between brain volumes and 57 major diseases using brain MRI scans in UKB, but their approach was biased by a limited number of pre-selected diseases and manually engineered features.

Thus, a scalable framework to systematically uncover associations between image-derived features and a broad range of diseases is still lacking. Such a disease-wide approach would be less constrained by prior assumptions or domain-specific biases, enabling the discovery of previously unrecognized patterns and potential biomarkers. This could enable improvements in incidental diagnosing capabilities, potentially leading to earlier detection of diseases and improvements in quality of life, as well as decreased healthcare utilization. For example, incidental detection of hypothyroidism can allow the prevention of pericardial effusion and the potential progression to cardiac tamponade (Kaur et al. 2021). Screening of Splenic Gaucheroma in CT scan images can incidentally uncover previously unrecognized Gaucher disease, enabling earlier intervention (Erdal et al. 2023). Incidental pulmonary embolism was detected early on routine cancer-staging chest CT scans, allowing rapid treatment before symptoms developed (Banala, Srinivas R. et al. 2017).

To address this gap, we developed an SSL-based framework, termed image-based disease-wide association study (iDWAS), to derive features from abdominal MRI images and evaluate their associations with a range of diseases in UKB **(Fig. 1)**. Our framework revealed multiple significant associations between the MRI features and various diseases. We specifically examined the association with MASH in depth and observed that MRI-derived features could perform better than the widely used PDFF (Boncan et al., 2024) and iron-cT1 (Dennis et al., 2021) biomarkers in separating MASH cases and controls.

**Fig. 1.**
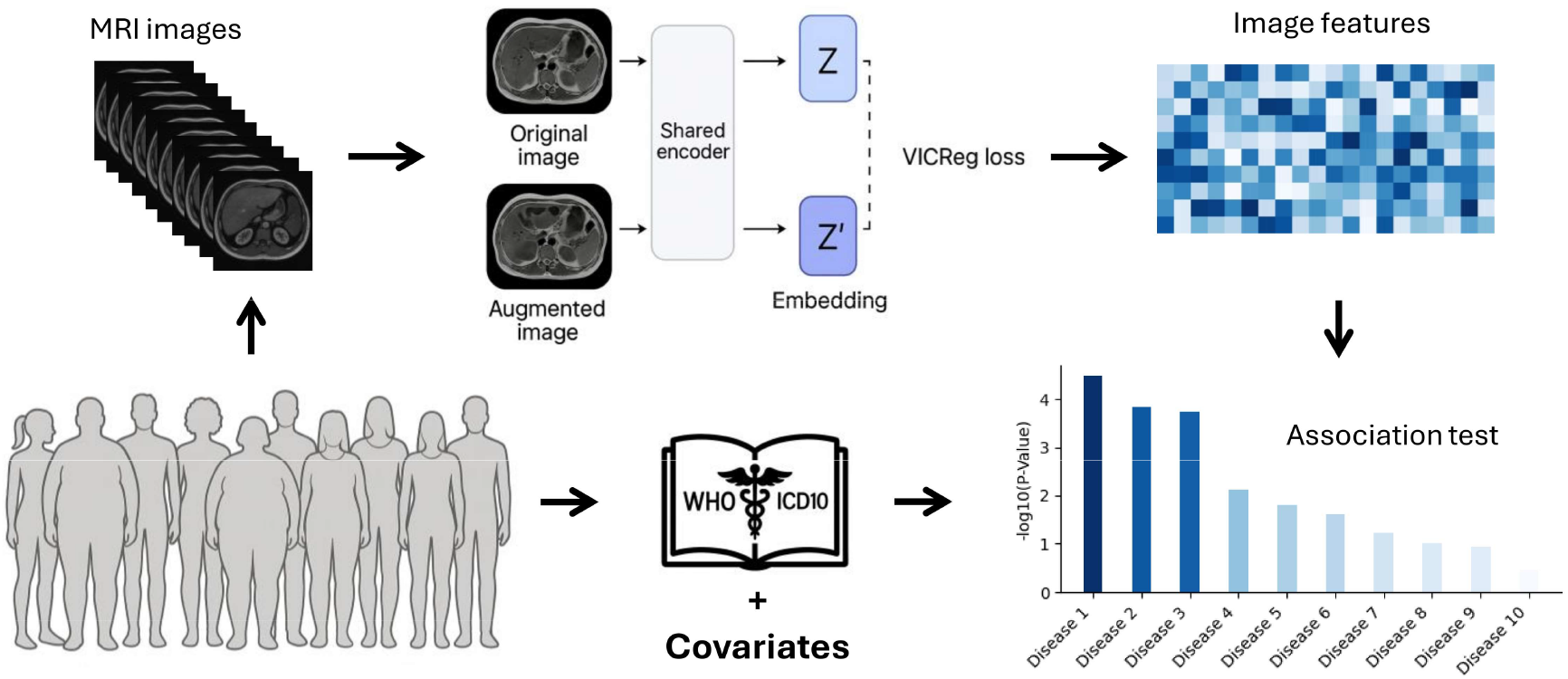
Schematic representation of the Image-based disease-wide association study (iDWAS) framework.

## Methods

### Data and cohort

We downloaded the UKB data from DNANexus^1^ using its Command Line Interface (CLI) dx-toolkit^2^. From a total of 502,170, we selected only those 60,662 participants for whom abdomen MRI scans of IDEAL protocol, more specifically with nonempty field p20254_i2, were available. These participants were re-contacted from the original UKB cohort, with selection based on willingness to participate and influenced largely by proximity to imaging centres (e.g., travel distance).

We further filtered the participants for self-reported white ethnicity to reduce confounding from population stratification, i.e. those with ‘British’, ‘Irish’, ‘Any other white background’, or ‘White’ values in their p21000_i0 field, resulting in 58,459 participants.

Of these 58,459 participants, 49,865 (85.3%) were diagnosed with at least one disease, representing a total of 7204 distinct diseases. **Fig. S2** shows the number of diseases per person and the prevalence of the 15 most frequent diseases.

### Feature extraction from MRI Images

As annotated data for abdominal MRI images is limited and costly to obtain, we adopted a self-supervised learning (SSL) framework to extract robust image features that can be used in downstream tasks **(Fig 1 and S1)**. Specifically, we used VicReg (Variance-Invariance-Covariance Regularization) [Bardes et al., 2022] to embed the images in a 512-dimensional feature space. As the backbone, we employed a ResNet-34 architecture [He et al., 2016], adapting its first convolutional layer to handle grayscale MRI inputs and removing its final classification layer. The learned representations were projected using a four-layer perceptron with dimensions 512 → 128 → 128 → 128. The VicReg loss function combines three complementary terms, weighted by coefficients of 25 for variance, 25 for invariance, and 1 for covariance. For the training, we generated two augmented views of each input image by applying a sequence of random transformations, including colour jitter to modify brightness and contrast by ±10%, random rotation of up to ±5°, random affine transformations shifting the image horizontally and vertically by up to 3% and 2%, respectively, and random resized cropping of 90–100% of the original size followed by resizing to 224×224 pixels. To improve robustness, we also added Gaussian noise (mean = 0, standard deviation = 0.001) and normalized pixel intensities using a mean and standard deviation of 0.5. The model was trained using standard stochastic gradient descent (SGD) with a learning rate of 0.01, a batch size of 256, and for 1,000 epochs. The choice of 1,000 training epochs was guided by empirical observations during model development. In preliminary experiments with shorter training schedules (e.g., 100–200 epochs), the learned representations had not yet stabilized, whereas extending training to 1000 epochs consistently resulted in stable embeddings, with no further improvements seen beyond this. We did not use a separate validation set or early stopping criterion during self-supervised training, as VICReg-based approaches are generally less prone to overfitting due to their self-supervised nature (no label-overfitting) and regularization mechanisms (variance, invariance, and covariance terms). Instead, we relied on the stability of the learned representations and downstream task performance as an implicit validation.

### Disease dictionary and data cohort

We termed any medical condition for which an ICD10 (International Classification of Diseases 10th Revision) code exist(s) (WHO, 2016; WHO, 2019) as a disease in this study. We considered version 2019 of ICD10-UK codes (here onwards referred as ICD-10 codes). All diseases assigned to the UKB participants in our cohort, their ancestors (i.e. their containing categories) and all the non-injury non-ill-defined secondary-level mortality causes defined by World Health Organization (WHO) were included in our disease dictionary. The former were collected from the *Diagnoses-ICD10* (p41270), *Underlying (primary) cause of death ICD10* (p40001_i0), and *Contributory (secondary) causes of death ICD10* (p40002_i0_a0 to p40002_i0_a9) data fields, and the latter from WHO (2025).

We defined a proxy-MASH cohort from the UKB participants using a combination of ICD10-based inclusion and exclusion criteria, based on Vujkovic et al. 2022 **(Table S3)**.

### Association between individual features and UKB variables

We used the linear regression models *UKB variable ∼ MRI-derived feature* to test for the associations of individual features with UKB variables namely Age, Gender, liver iron corrected T1 (Iron-cT1), Liver_volume, Body Mass Index (BMI), Proton Density Fat (PDFF), Alanine Aminotransferase (ALT), Aspartate Aminotransferase (AST), Gamma Glutamyl Transferase (GGT), and the first five genetic Principal Components (GPC_1_ to GPC_5_).

The UKB variables, except for Gender and the imaging features, were normalized before the analysis. In addition to the UKB variables, we randomly synthesized six ‘dummy’ variables: three continuous variables drawn from the standard normal distribution and three binary variables (coded as ‘0’ or ‘1’) drawn from the Bernoulli distribution with three different probability values for the label ‘1’ as 0.5, 0.1, and 0.01. For each model, the P-value and the effect size (i.e. the absolute value of the coefficient) were reported.

### Association between MRI-derived features and diseases, including proxy-MASH

We observed 7204 distinct disease codes assigned to patients in the UKB database, of which 6249 were among valid 2019 edition of ICD10-UK codes (here onwards referred as ICD-10 codes), and the rest (955) were skipped. The number of distinct ancestors of the included diseases was 1813. These ancestors include chapter numbers, which are still referred to as ‘disease’ here, for simplicity. For example, the ancestors of the ICD10 code ‘K76.81’ are ‘K76.8’, ‘K76’, ‘K70-K77’, and ‘XI’, where the last one is a chapter number. In addition, we extracted from WHO 170 causes of death. For simplicity, we also refer to these causes of death as ‘disease’ even though many of them contain one or more ranges of ICD10 codes. Of these 170 diseases, 73 had already been among the identified UKB diseases or their ancestors, and the remaining 97 were added to the selected list of diseases. These diseases together with our proxy MASH definition, make a total number of 8160 diseases, among which there were 562 diseases with at least 500 cases.

Considering only these 562 diseases with at least 500 cases, we used the following regression models to test for associations between the individual features and each disease, while adjusting for potential confounding by age, gender, BMI, and first 5 genetic PCs. There might be unobserved noise in the labels is this public dataset, which could not be accounted for.

Base model:

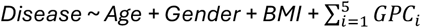

Full model:

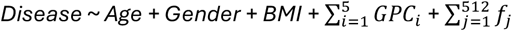

where variable i refers to the number of Genetic PCs (GPCs) and variable j refers to the number of MRI-derived features (embeddings).

We evaluated the full model against the base model using the Likelihood Ratio (LR) test. The relative reduction in the log loss, a measure of improvement in the model fit due to including MRI-derived features was calculated as (*ll_base – ll_full / ll_base*, where *ll_base* and *ll_full* are the log loss values of the individual base and full models, respectively.

For proxy-MASH specifically, we additionally adjusted for potential confounding by smoking and alcohol usage as follows:

Base model:

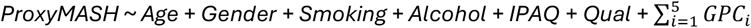

Full model:

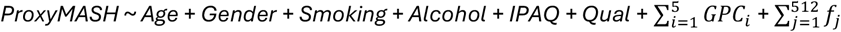

where IPAQ refers to International Physical Activity Questionnaire and Qual refers to educational qualification.

We evaluated MASH biomarkers/outcomes PDFF and Iron-cT1 as follows:

Base model:

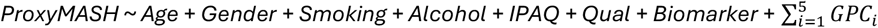

Full model:

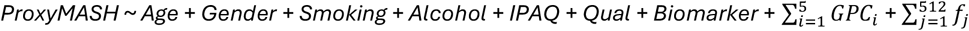

*where Biomarker would be PDFF or/and Iron CT1*.

The associations of the imaging features with the diseases (including MASH) were also examined after feature selection. We used the feature selection method in Tegtmeyer, Arora et al. (2024), selecting uncorrelated features by requiring each pair of selected features to show a correlation coefficient less than a specific threshold. We evaluated thresholds ranging from 0.1 to 0.95 with step size 0.05.

## Results

First, we report on the associations of the individual MRI-derived features with common confounders and/or liver markers in UKB. Then, we investigate the association of the combined set of MRI-derived features across the disease spectrum, with a specific focus on MASH.

### MRI-derived features demonstrate biological relevance

We first assessed the associations of each of the 512 MRI-derived features with common confounders such as age, gender, BMI, and liver volume, as well as with liver related biomarkers such as PDFF, Iron-cT1, ALT, AST, and GGT (**Fig. 2**). Among these, gender showed the strongest associations, suggesting that the MRI-derived features captured gender-specific differences in the abdomen. GPCs showed the weakest associations, potentially due to the inclusion of individuals with white ancestry only, leaving to relatively genetically homogeneous cohort. We confirmed these observations with a permutation-based analysis **(Fig. S3)**.

**Fig. 2.**
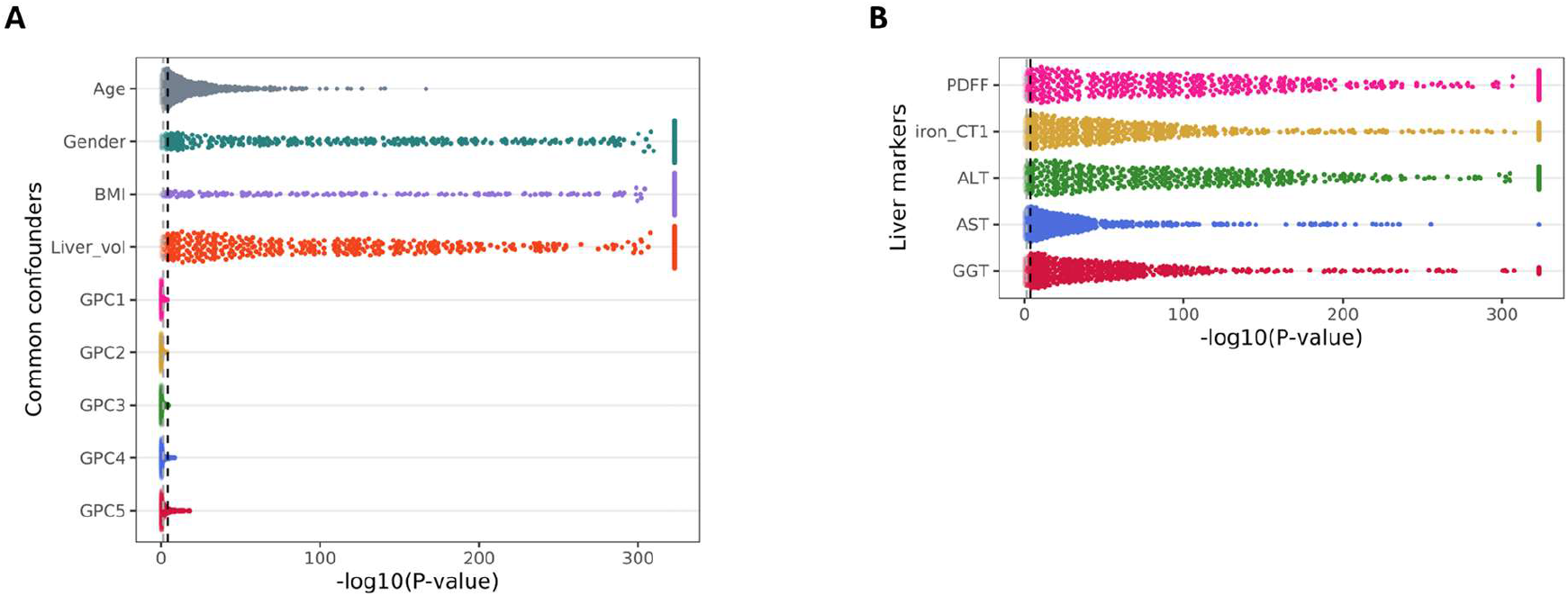
Associations of MRI-derived features with UKB variables. Associations of each of the 512 MRI-derived features with common confounders **(A)** and liver biomarkers **(B)** in UKB, where each feature is represented by a single point. The x-axis is capped at 323 which corresponds to the smallest nonzero P-value that can be represented in a standard Python variable. The grey and black dotted vertical lines represent P-value thresholds of 0.05 and 9.7×10^-5^, respectively, the latter representing the Bonferroni-corrected P-value threshold (P < 0.05 / 512).

### Association of MRI-derived features with multiple diseases

As studies have shown that ML-based analyses of MRI images can capture more disease-relevant information than human observers or standard biomarkers, so we hypothesized that the SSL-based features from abdominal MRI images potentially capture information relevant for medical conditions. To test our hypothesis, we assessed the association of MRI-derived features with the diseases defined with ICD10 codes by WHO (see Methods).

Out of a total 8160 diseases in the final set, we selected the 562 diseases presented with at least 500 cases (see Methods, **Fig. S2**). We then regressed case-control status of each of these 562 diseases on the 512 MRI-derived features and observed that the MRI-derived features explained significantly more variation in case-control status in 158 diseases, compared to considering only baseline covariates age, gender, BMI and genetic PCs (LR test *P*-value < 0.05/562 = 8.9×10^-5^, **Fig. 3 and Fig. S7, S8**).

**Fig. 3.**
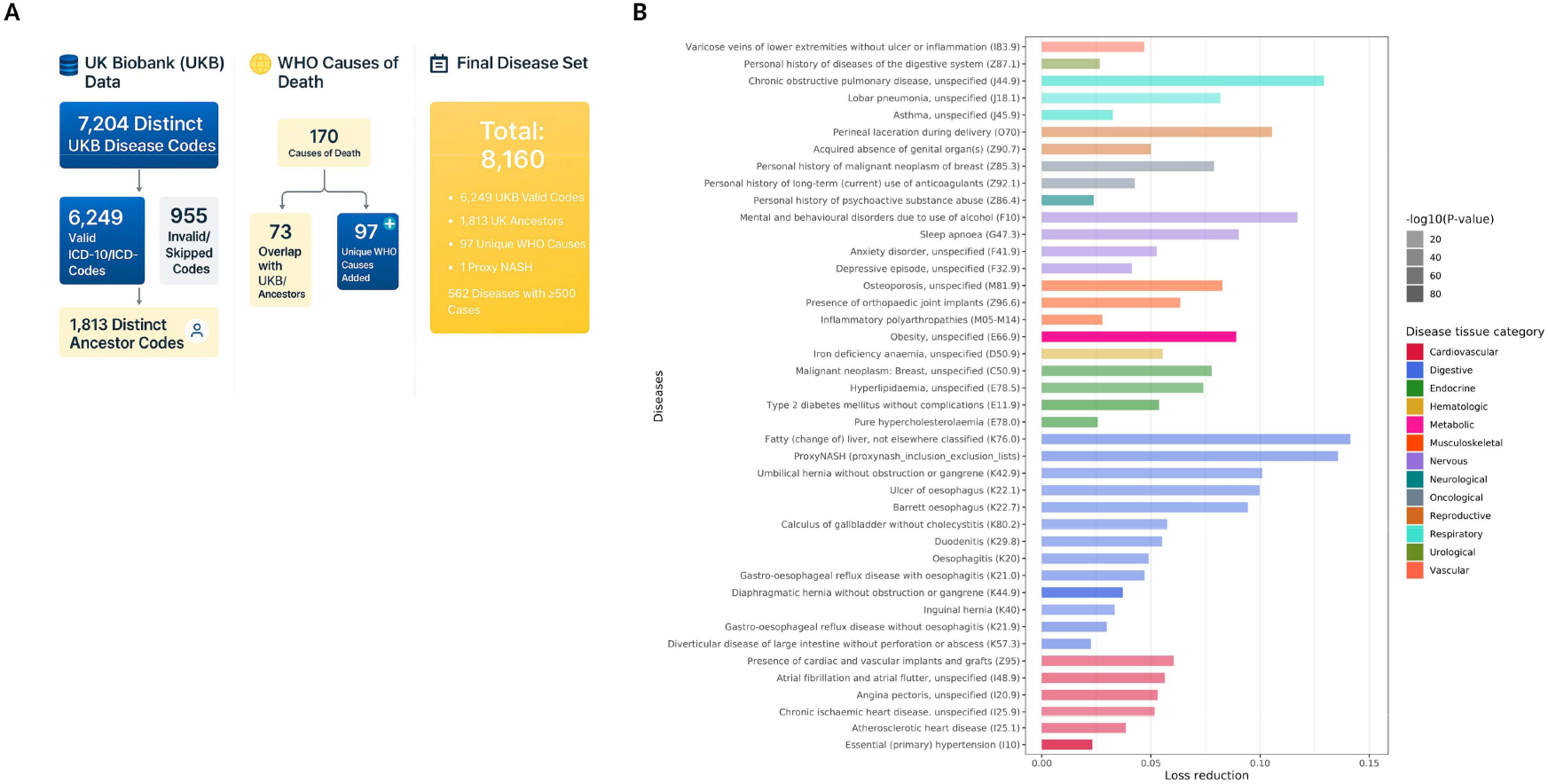
Image-based disease-wide association study (iDWAS) **(A)** Harmonization of ICD10 disease codes and cohort selection from UKB for downstream association analysis. **(B)** Diseases, grouped by their anatomical site, for which the MRI features explained case-control status significantly better than baseline covariates are shown. Loss reduction represents the relative improvement of the model due to including the MRI-derived features over baseline covariates only.

Among these 158 diseases, we observed many diseases which primarily emerge in the abdomen. For example, type 2 diabetes mellitus (ICD10 code E11) is linked to visceral adiposity and hepatic steatosis, both detectable via MRI scans (Bril & Cusi, 2016). Gastro-oesophageal reflux disease (K21) might reflect diaphragmatic or stomach changes, such as hiatal hernia or increased intra-abdominal pressure (Zhou et al., 2022). Fatty liver disease (K76.0) is also directly measurable on MRI scan, which is sensitive to hepatic fat content (Caussy et al., 2018).

Interestingly, among the 158 diseases, we also observed diseases not traditionally directly linked to abdominal anatomy **(Fig. 3)**, such as sleep apnoea (G47.3) and depressive disorders (F32). Both disorders are however known to associate with systemic inflammation and visceral adiposity (Shinohara et al., 2020, Lee et al., 2012; Zhao et al., 2022), which could be captured in abdominal MRI. Another example not traditionally linked to abdominal anatomy is osteoporosis (M81.9). Increased bone marrow fat, captured via MRI, has however been shown to inversely correlate with bone mineral density in adult cohorts (Shen et al., 2012), and moreover marrow adiposity is increased in osteoporosis (Justesen et al., 2001). These findings illustrate that the features derived from abdominal MRI images using our SSL-based framework can capture not only abdomen-specific pathology but also aspects of systemic disease processes occurring potentially distal to the abdomen.

### Segregation of MASH cases and controls

Next, we analyzed the associations of our MRI features with Metabolic dysfunction–associated steatohepatitis (MASH) in more depth. MASH is the most severe form of Metabolic dysfunction-associated Fatty Liver Disease (MAFLD), characterized by excessive hepatic fat accumulation along with inflammation and hepatocellular injury. It is estimated to affect approximately 3–6% of the global adult population (Povsic et al. 2019), with U.S. estimates ranging between 3% and 5% (Estes et al. 2018b, Younossi et al. 2019, Rinella 2015). MASH imposes a significant burden on both individuals and society due to its potential progression to cirrhosis, hepatocellular carcinoma and liver failure (Estes et al., 2018a) and is associated with increased cardiovascular risk and reduced quality of life (Targher et al., 2010).

Due to the unspecific nature, or even absence, of early-stage MASH symptoms, MASH remains substantially under-diagnosed (Schattenberg et al., 2023; Fialoke et al., 2018). Hence, we used a combination of ICD10 codes, termed as proxy-MASH, to define a cohort of 561 individuals with a MASH-like profile while excluding for known differential diagnoses (see Methods, **Table S3** and **Fig. S4** for MRI examples of proxy-MASH patients and a visualization of the derived features). Next, we evaluated how well MRI-derived features could separate the proxy-MASH cases from the remaining individuals and observed that the MRI-derived features explained significantly more variation in proxy-MASH status compared to considering only the baseline covariates age, gender, smoking, alcohol, IPAQ and qual (Likelihood Ratio (LR) test *P* = 2.4 × 10^-25^, Loss Reduction = 0.19, **Fig. 4A**).

**Fig. 4.**
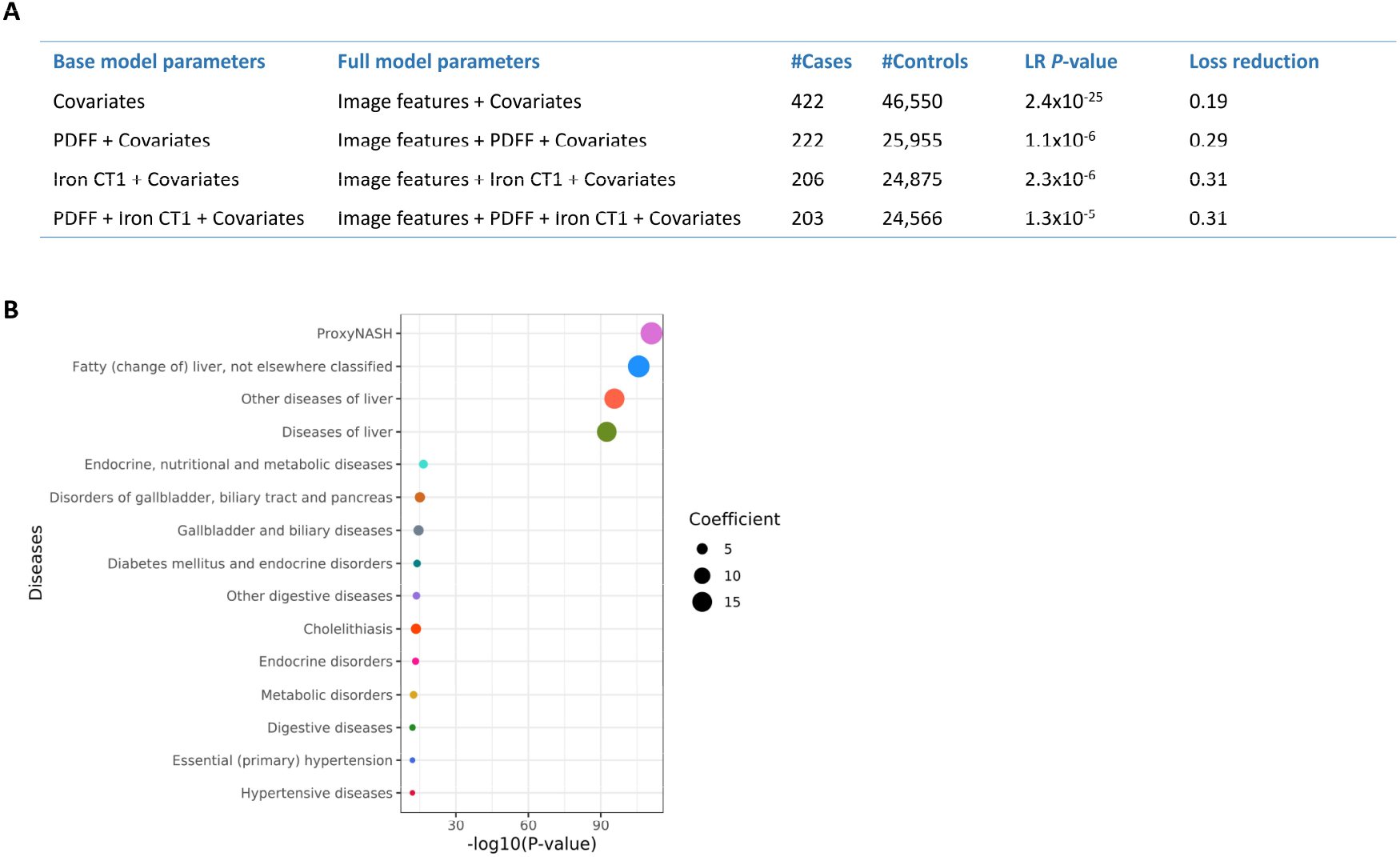
Association of the MRI-derived features with proxy-MASH. **(A)** The table shows the improvement in separating proxy-MASH cases from controls by MRI-derived features over the covariates, PDFF and Iron-CT1. Loss reduction corresponds to the improvement in the fit of full model over base model. **(B)** Diseases with significant associations with the difference (Δp) in fitted probabilities between the proxy-MASH models with and without 512 MRI-derived features (using Bonferroni-corrected P-value threshold (P < 0.05 / 562)). Only top 15 diseases out of 77 significantly associated ones (Fig. S9) are shown here.

We then asked whether the MRI-derived features could explain more variation in proxy-MASH status than the widely used MASH biomarkers of Proton Density Fat Fraction (PDFF) and Iron-corrected T1 (Iron-cT1) (Boncan et al., 2024; Dennis et al., 2021), for which 222 and 206 proxy-MASH cases were available in UKB, respectively. We observed that adding the MRI-derived features to models containing either only PDFF or only Iron-cT1 significantly improved the fit of the model **(Fig. 4A)**. Thus, the MRI-derived features capture information-relevant information from the MRIs that is not currently captured by standard MASH biomarkers. Notably, a model including both PDFF and Iron-cT1 only marginally improved the model fit compared to Iron-cT1 alone.

To ensure that these findings were not an artifact of sample size, we confirmed the findings using a re-sampling approach by randomly selecting 222 and 206 participants with PDFF and Iron-cT1 information as proxy-MASH cases and repeating the above analysis 1000 times to calculate a background distribution of LR *P*-values (**Fig. S6**). To ensure that the above findings were not an artefact of the relatively high dimensionality of the model (N=512 MRI-derived features), we confirmed the findings in a number of lower-dimensional models, each based on a limited set of features selected by capping the inter-feature correlation (see Methods; **Fig. S6**). These tests strongly supported the statistical and biological relevance of our SSL-based framework and MRI-derived features. Moreover, we observed that features derived from the whole abdomen accounted for larger variation in MASH status compared to the features derived from only the liver. This is in alignment with a previous study by Pillai et al. (2022), where the whole abdomen provided a more accurate prediction of Non-Alcoholic Fatty Liver Disease (NAFLD) than liver only. Overall, these results demonstrate that the MRI-derived features capture information not captured by currently widely used MASH biomarkers, and that focusing on the whole abdomen can be more informative than focusing on the liver only.

Having observed that the MRI-derived features significantly improved the fit over PDFF and Iron-cT1, we wanted to gain more insight into the underlying reasons for the improvement in fit. For that, we considered the difference (Δp) in fitted probabilities between the models with and without the 512 image features as a measure of the additional variance in MASH status captured by the model that included the MRI-derived features, relative to the model that did not include those features. We then calculated the association of Δp with disease status for every disease covered in Section 3.2 (see Methods). We identified significant associations for 77 out of 562 diseases **(Fig. 4B)**, highlighting that the additional pathological information captured by the MRI-derived features can relate not only to the abdomen itself, but also co-relate to indications typically associated with organs outside the abdomen, which may not be captured by PDFF and Iron-cT1 alone. An interesting example here is a neurological disorder such as migraine, known to be associated with NAFLD (Celikbilek, et al., 2014), whose most severe form is MASH. In particular, Celikbilek et al. showed that among migraine patients, those with NAFLD had significantly higher measurement of BMI, waist circumference and multiple biomarkers including insulin levels ALT and triglycerides in comparison to those without NAFLD.

## Discussion

In this study, we applied SSL on abdominal MRI scans in the UKB to obtain application-agnostic features. We used these features as predictors in an image-based disease-wide association study (iDWAS) and showed significant association with 158 diseases. Besides the expected associations, such as with MASH, our features associated with various diseases occurring in tissues other than the abdomen, such as neuropsychiatric diseases. Notably, the MRI-derived features captured additional variation in proxy-MASH case and control status compared to the widely used PDFF and iron-cT1 biomarkers.

### Study limitations

While the present study demonstrates the utility of self-supervised learning (SSL) for extracting clinically relevant features from abdominal MRI data, several limitations of this study should be acknowledged, while simultaneously highlighting opportunities for further methodological and biological refinement. First, the SSL model was trained without the use of a dedicated validation set or early stopping criterion. Although VICReg-based frameworks incorporate regularization mechanisms that may mitigate overfitting, future work could benefit from explicitly incorporating validation strategies to enable more systematic assessment of generalization performance and optimization of training procedures. Second, the imaging pipeline was intentionally simplified by relying on a single two-dimensional MRI slice per participant. While this approach ensures consistency and computational efficiency, it presents an opportunity to extend the framework to multi-slice or fully three-dimensional representations, which could better capture the spatial complexity and heterogeneity of abdominal anatomy and pathology. In addition, using a single MRI image per donor, combined with the cross-sectional nature of the study, does not account for temporal discrepancies between imaging, clinical measurements, and disease diagnoses. Third, disease labels derived from UK Biobank data are subject to potential misclassification. For example, participants without a recorded ICD-10 code for a given disease were treated as controls, which may result in inclusion of undiagnosed or preclinical cases in the control group. In addition, the Proxy-MASH phenotype is itself subject to residual misclassification, as it relies on surrogate diagnostic criteria rather than histologically confirmed disease status. More broadly, unobserved label noise inherent in large public datasets may affect the robustness of association results. Fourth, the imaging cohort was derived from UK Biobank participants who underwent abdominal MRI as part of the imaging extension study, which relied on re-contacting and recruiting participants from the original cohort. As a result, the imaging sample may be affected by selection bias and healthy-volunteer bias, with participants undergoing imaging potentially being healthier and more engaged than the broader population. Furthermore, the study population was restricted to individuals with self-reported white ethnicity to reduce confounding from population stratification. While this improves internal validity, it limits the generalizability of the findings to more diverse populations. Fifth, although the learned MRI-derived features capture disease-relevant information, their biological interpretation remains an open area for investigation. In particular, integrating imaging-derived representations with genetic data (e.g., through genome-wide association studies) offers a promising avenue to uncover the molecular and genetic underpinnings of the extracted features and to enhance mechanistic insight. Finally, several modeling decisions in the current pipeline, such as the choice of embedding dimensionality (fixed at 512) and other architectural or training hyperparameters, were not systematically explored. This presents an opportunity for future work to perform more comprehensive model selection and sensitivity analyses, which may further improve predictive performance and robustness.

### Conclusion

Collectively, the above limitations outline a set of concrete opportunities to extend the present framework, including improved validation strategies, richer imaging representations, more refined phenotyping, and integration of multi-modal data. In conclusion, we demonstrated through an abdominal MRI use case that SSL can enhance information extraction from MRI data compared to standard clinical biomarkers. Additionally, we showed that liver MRI-derived features can have cross-organ relevance. Combined, we believe that these results illustrate that the data and tools are now available to begin to think systematically about improving incidental diagnosing capabilities, thus opening opportunities for earlier and better detection of diseases, and ultimately improving quality of life.

## Supporting information

All Supplementary

## Acknowledgements

This work performed at Glasgow University was funded by Boehringer Ingelheim Pharma GmbH. This research has been conducted using the UK Biobank Resource under Application Number 71392. We are very thankful to all participants in the UK Biobank and the project team. We also thank Sina Mousavi for insights on incidental diagnoses using available ICD-10 codes in UKB, as well as contributions to disease-wide association studies and the anatomical characterization of the diseases included in this study.

## Data availability

The source code and the trained model are publicly available at https://github.com/srm2022/iDWAS.

## Footnotes

1 https://ukbiobank.dnanexus.com

2 https://documentation.dnanexus.com/getting-started/cli-quickstart.

## Notes

### Competing Interest Statement

The authors have declared no competing interest.

