## Supplementary material for "Image-based Disease-wide Association Study via Self-supervised Learning links Abdominal MRI Features to 158 Diseases": All Supplementary

### A. Supplementary materials related to methods

#### A.1. Manual and automated modules in the pipeline

**Table S1. Summary of the pipeline's modules, together with their inputs and outputs.** Table summarises the manual (UKB1 and UKB2) and automated (IMG, SSL, WO, ICD, VAR, iDWAS, and MASH) modules in the pipeline, and Fig. S.1 illustrates their interconnections.

| Name | Type | Short description | Input data | Output data | Program files |
|---|---|---|---|---|---|
| UKB1 | Manual | To get UKB MRI zip files | UKB data via DNANexus | A folder containing the IDEAL-protocol MRI scan zip files | N/A |
| UKB2 | Manual | To get health-record data | UKB data via DNANexus | data_participant.csv | N/A |
| IMG | Automated | To create one png file per each MRI scan zip file | UKB1-out | A folder, e.g. 'all_images', containing png files | pipe_img.ipynb |
| SSL | Automated | To train an SSL model on images | IMG-out | A JSON files containing the H features of the input images | pipe_ssl.ipynb |
| WHO | Automated | To get disease names and icd10 codes from WHO | Names and lists of ICD10 codes of diseases in WHO | who_icd10.tsv | pipe_who.ipynb |
| ICD | Automated | To identify cases for each disease | 1-UKB2-out<br>2-WHO-out | A pickle file disease_cases.pkl containing diseases' ICD10 codes together with the cases' EIDs | pip_icd.ipynb |
| VAR | Automates | To find associations for several UKB variables | 1- SSL-out<br>2- UKB2-out | Results | Pipe_VAR.ipynb<br>utils_ukb.py |
| iDWAS | Automated | To find associations for all diseases | 1- SSL-out<br>2- UKB2-out<br>3- ICD-out | Results | pipe_idwas.ipynb<br>utils_ukb.py<br>utils_stat.py |
| MASH | Automated | To find and analyse associations for MASH | 1- SSL-out<br>2- UKB2-out<br>3- ICD-out (only for proxy-MASH) | Results | pipe_mash_no_pdff.ipynb<br>pipe_mash_pdff.ipynb<br>pipe_mash_iron_ct1.ipynb<br>pipe_mash_both_pdff_iron_ct1.ipynb<br>pipe_mash_diff_analysis.ipynb<br>utils_ukb.py<br>utils_stat.py |

**Fig. S1. Interconnections among the modules in the pipeline**

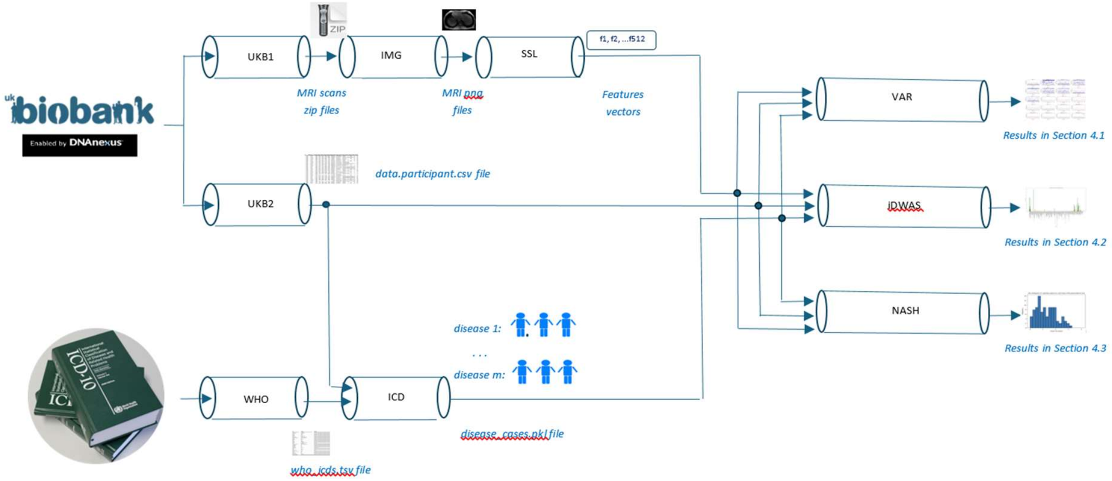


### Manual modules

The manual modules UKB1 and UKB2 download the image and health-record data, respectively, from UKB and produce the following:

- A folder containing zip files, one zip file per participant
- A csv file containing the health record data, one row per participant

### Automated modules

The rest of the modules in Table S.1 are automated. The WHO module gets ICD10 codes of diseases from a specific WHO webpage[1]. This module is hard-coded, which means that it will need to be modified if the content and/or the structure of this webpage is modified. The remaining modules consist of two groups:

- Group 1: IMG, SSL, and ICD.

  IMG generates a png image file per participant. SSL trains a VICReg SSL model and generates a dictionary that contains one feature vector per participant. ICD generates a dictionary that contains for each ICD10 code (or a group of ICD10 codes) the corresponding disease name and list of the cases' EIDs.

- Group 2: VAR, iDWAS, and MASH.

  These modules only perform the association analyses and generate the data of the graphs and tables presented in the paper. All independent variables in the regression analyses are normalised.

Further notes about the above manual and automated modules are provided next.

[1] https://platform.who.int/mortality/about/list-of-causes-and-corresponding-icd-10-codes

### UKB1

The filenames of the MRI scans were obtained from the p20254_i0 field. The dx-toolkit was used to find and download the images. It could find and download 59427 out of 60662 files.

### UKB2

The generated csv file contains the data fields shown in Table S.2 (in addition to the Encoded Identifier, EID).

**Table S2. UKB data fields selected by the UKB2 module.**

| **Field ID** | **Field name** | **Name in our analyses** | **Values in our analyses** |
|---|---|---|---|
| p21003_i0 | age attended assessment centre | *Age* | (no change) |
| p31 | sex | *Gender* | Female: 0; Male: 1 |
| p21001_i0 | Body Mass Index | *BMI* | (no change) |
| p22009_a1 to p22009_a5 | The first 5 genetic Principal Components | $GPC_1$ *to* $GPC_5$ | (no change) |
| p20116_i0 | smoking status | *Smoking* | Never: 0; Previous: 1; Current: 2 |
| p1558_i0 | alcohol intake frequency | *Alcohol* | Never: 0<br>Special occasions only: 1<br>One to three times a month: 2<br>Once or twice a week: 3<br>Three or four times a week: 4<br>Daily or almost daily: 5 |
| p22032_i0 | IPAQ activity | *Ipaq* | Low: 1; Moderate: 2; High: 3 |
| p6138_i0 | Qualifications | *Qual* | Other professional qualifications e.g: nursing, teaching: 6<br>College or University degree: 5<br>A levels/AS levels or equivalent: 4<br>O levels/GCSEs or equivalent: 3<br>CSEs or equivalent: 2<br>NVQ or HND or HNC or equivalent: 1<br>None of the above: 0 |
| p40061_i2 | Proton Density Fat Fraction | *PDFF* | (no change) |
| p40062_i2 | Liver iron corrected T1 (cT1) | *Iron_CT1* | (no change) |
| p21080_i2 | Liver volume | *Liver_volume* | (no change) |
| p30620_i0 | Alanine Aminotransferase | *ALT* | (no change) |
| p30650_i0 | Aspartate Aminotransferase | *AST* | (no change) |
| p30730_i0 | Gamma glutamyltransferase | *GGT* | (no change) |
| p41270 | Diagnoses - ICD10 | N/A | N/A |
| p40001_i0 | Underlying (primary) cause of death ICD10- Instance 0 | N/A | N/A |
| p40002_i0_a0 to p40002_i0_a | Contributory (secondary) causes of death ICD10- Instance 0- Arrays 0 to 9 | N/A | N/A |

UK Biobank education categories are not intrinsically ordered; therefore, we constructed an ordinal scale based on functional educational attainment. We ranked 'other professional qualifications' above 'college or university degree' because they can represent regulated, post-academic credentials that reflect higher occupational specialization and labour-market readiness than a degree alone, such as medical doctor, accountant, lawyer. We do however recognize that this category is heterogeneous and can also include mid-level certifications.

#### IMG

It receives as input the output of UKB1 (i.e. a folder containing the MRI zip files) and outputs a single png file for each zip file. Three zip files could not be opened, and one was skipped because of containing images of 256 x 232 pixels (as opposed to 232 x 256 of all other images), resulting in 59423 png files, stored in a folder called 'all_images'. Each zip file contained 72 images in the DICOM format for a participant, 36 magnitude and 36 phase images. We consistently selected the magnitude image with Acquisition Number 1 and Instance Number (echo time) 5 for all participants. These numbers are parts of the metadata in the DICOM files. This Instance Number was selected because it corresponds to an echo time (9.2 ms) at which signals from fat and water are in phase. Under this condition, their signals add constructively, resulting in a higher and more stable signal-to-noise ratio while avoiding systematic signal cancellation artifacts. By reducing phase-dependent intensity variations and chemical shift-related artefacts, the respective images provide a more homogeneous input distribution. This minimizes acquisition-dependant signal fluctuations and supports the model encoding meaningful pathological or anatomical features rather than non-biological, phase-related artefacts.

#### SSL

We train a VICReg model on PNG images (received from the IMG module) and produce as output their features, saved as a JSON file. For each input image $t_1$, two stochastic augmentations $t_2$and $x_1 = t_1(x)$ were sampled to generate two correlated views $x_2 = t_2(x)$ and $f_\theta(.)$. Both views were processed by a shared encoder $g_\varphi(.)$ followed by a projection head $z_1 = g_\varphi(f_\theta(x_1))$, producing projected embeddings $z_1 = g_\varphi(f_\theta(x_1))$ and $t_1$. The augmentations $t_2$and $f_\theta$ were generated using random transformations: brightness/contrast jitter of ±10%, random rotation up to ±5°, random affine translation up to 3% horizontally and 2% vertically, and random resized cropping retaining 90–100% of the original area followed by resizing to 224 × 224 pixels. Gaussian noise (mean 0, standard deviation 0.001) was added and pixel intensities were normalised using mean 0.5 and standard deviation 0.5. The encoder backbone $g_\varphi$ was a ResNet-34 adapted for grayscale MRI by modifying the first convolution to accept a single input channel and removing the final classification layer. Encoder outputs were 512-dimensional. The projection head was a four-layer multilayer perceptron (MLP), where the first layer had 512 neurons and the others 128 each.

VICReg optimizes a weighted sum of three terms: (i) an invariance term $L_{inv}$ that encourages view-consistent representations by minimizing the mean squared error between paired embeddings; (ii) a variance regularization term $L_{var}$ that enforces a minimum per-dimension standard deviation across the batch to prevent representational collapse; and (iii) a covariance regularization term $L_{cov}$ that penalizes off-diagonal entries of the batch covariance matrix, encouraging decorrelated (non-redundant) features. The total loss is $L = \lambda_{inv} L_{inv} + \lambda_{var} L_{var} + \lambda_{cov} L_{cov}$ with $(\lambda_{inv}, \lambda_{var}, \lambda_{cov}) = (25, 25, 1)$. The model was trained with stochastic gradient descent (learning rate 0.01, batch size 256) for 1,000 epochs. After training, each image was embedded using the encoder output (512-dimensional) as the feature representation; features were exported and stored in JSON format for downstream analysis.

#### WHO

It creates a csv file that contains the names and ICD10 codes of all the secondary-level causes of death defined by WHO (2025) except injuries and ill-defined diseases. The csv file called who_icd.tsv has three columns, one for disease name, one for (a concise representation of) the ICD10 codes (e.g. ‘A50-A53’), and one for the full list of the ICD10 codes (e.g. [A50, A50.1, ..., A52.9]). The first two columns were obtained from WHO (2025) and the last was generated by expanding the representations in the second column.

#### ICD

It receives as input the outputs of UKB2 and WHO and creates for each disease (including proxy-MASH), the list of EIDs of the participants diagnosed with the disease. That is, it obtains the cases for each disease. To this end, it first creates a dictionary of diseases from three sources: UKB, WHO, and our proxy-MASH ICD10 profile. From UKB, it imports every disease assigned to any participant in our cohort. It also uses WHO data indirectly (via two Python packages for ICD10 2019 and ICD10-cm 2019) to include the ancestors (i.e. super categories) of these diseases. Because the UKB disease names are shorts, they are replaced with those of ICD10 2019 / ICD10-cm 2019. From WHO mortality database, it imports indirectly those generated by the WHO pipe above. Finally, it uses the inclusion an exclusion list of ICD10 codes in Section A.3) to include proxy-MASH. The generated disease dictionary is stored in a dictionary, saved as a pickle file. Each key in the dictionary shows the ICD10 code(s) and each value is the pair of the associated disease name and EID list. It uses the inclusion and exclusion lists of ICD10 codes for proxy-MASH defined in Section A.3.

#### VAR

It generates the associations of each image feature with every UKB variable Age, Gender, Iron_CT1, Liver_volume, BMI, PDFF, ALT, AST, GGT, and the GPC1 to GPC5 as well as six dummy (random) variables, described in Section 4.1.

#### iDWAS

It obtains the associations between the whole set of the image features and every disease that has at least 500 cases among the participants. The covariates Age, Gender, BMI, and GPC1 to GPC5 are used in the logistic regression models.

#### MASH

It is like iDWAS except that it only finds associations for MASH, using more specialised covariates.

### A.3. Inclusion and exclusion lists of ICD10 codes for our proxy-MASH

Table S.3 shows the inclusion and exclusion lists of ICD10 codes used in-house to define proxy-MASH cases, increasing the number of cases from 25 (for K75.8) to 561.

**Table S3. The inclusion and exclusion lists of ICD10 codes for proxy-MASH defined in-house.**

| | |
|---|---|
| Inclusion list | K74.0, K74.00, K74.01, K74.02, K74.1, K74.2, K74.4, K74.5, K74.6, K74.60, K74.69, K75.8, K75.81, K76.0, K76.6 |
| Exclusion list | F10.9, F10.94, F10.95, F10.96, F10.97, F10.98, F10.99,<br>K70.0, K70.1, K70.2, K70.3, K70.4, K70.9, K75.4, K80.3, K83.0,<br>B18.0, B18.1, B18.2, B18.8, B18.9,<br>C22.0, C22.1, C22.2, C22.3, C22.4, C22.7, C22.8, C22.9,<br>E83.01, E83.11, E88.01 |

### B. Supplementary results

**Fig S2. Overview of diseases in participants from UKB who were included in this study.**

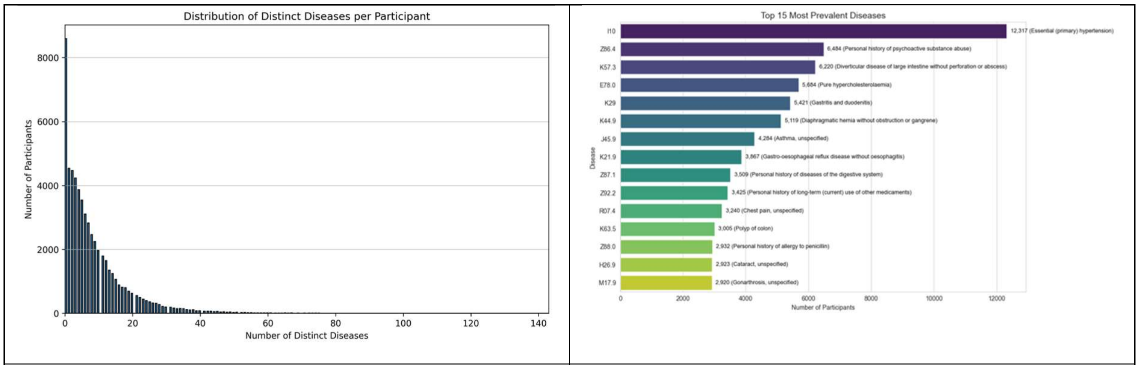

**Fig S3.** Associations of each of the 512 MRI-derived features (x-axis) with permuted confounding variables in UKB. The blue and red lines represent *P*-value thresholds of 0.05 and 9.7x10$^{-5}$, respectively, the latter representing the Bonferroni-corrected P-value threshold (*P* < 0.05 / 512).

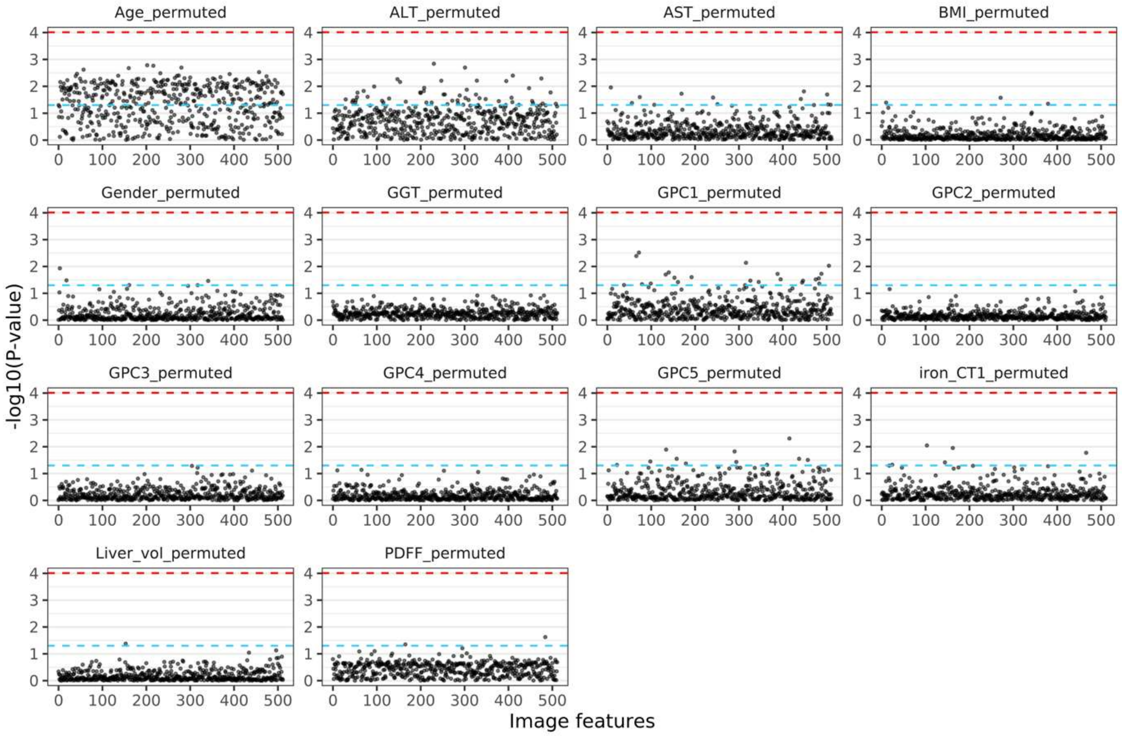


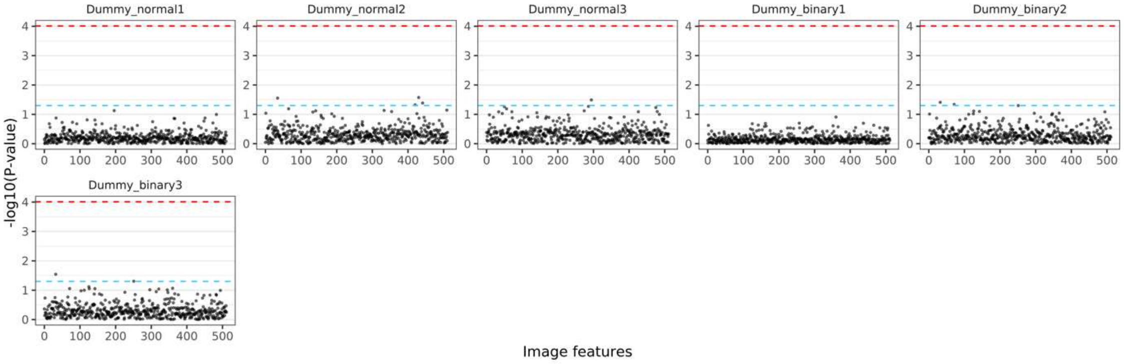

**Fig S4.** The first three MRI scans among those without MASH (left) and the first three among those with MASH (right) in our dataset.

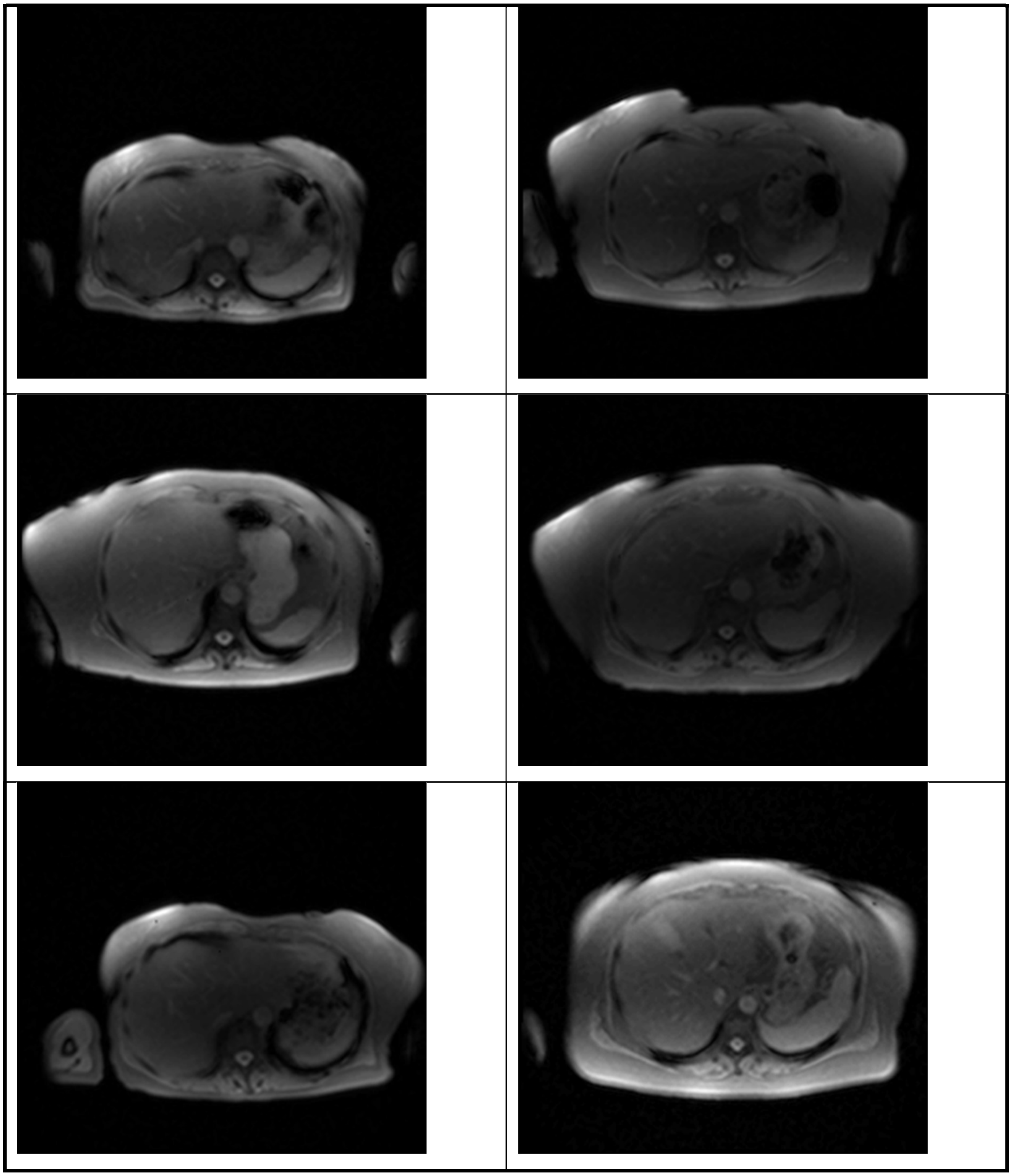

**Fig S5.** The 512-dimensional embeddings have been reduced to 2-dimension vectors using UMAP. The MASH and non-MASH instances have been shown in orange and blue, respectively. This simplified plot shows non-uniform distribution of the MASH cases, i.e. the MASH-relevance relevance of the features.

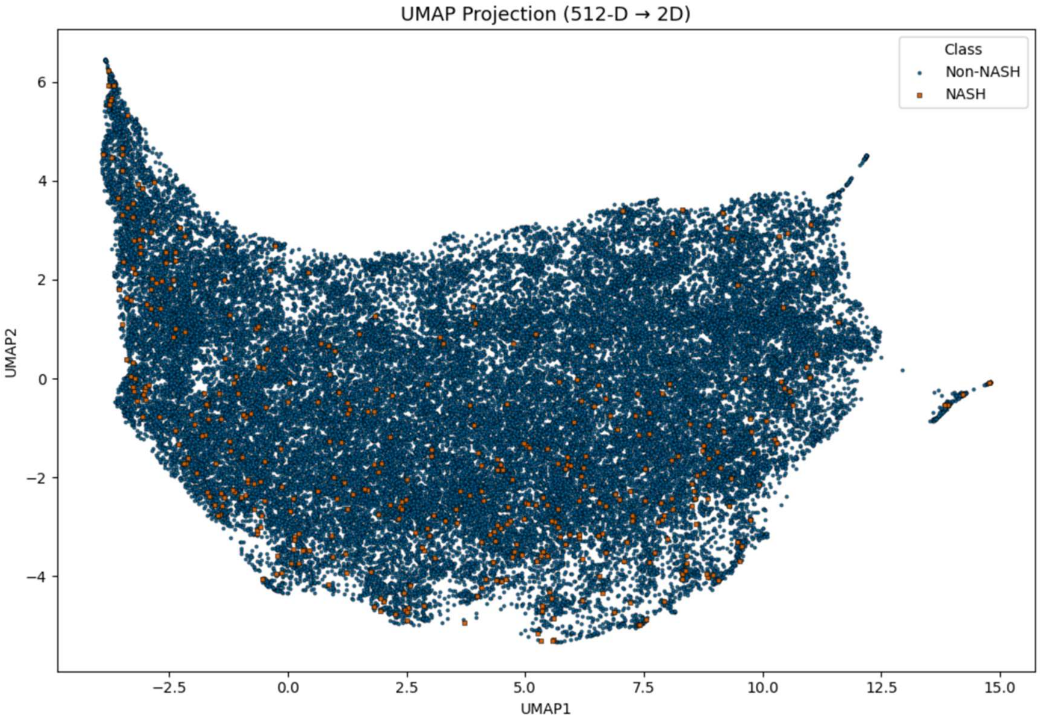


**Fig S6. (A)** Distribution of *P*-values obtained by random sampling of cases and controls from whole cohort for 1000 times showed that the observed significance (dotted lines) of additional R2 explained by features over PDFF and Iron-cT1 was not by chance. For PDFF, only 1 (out of 1000) *P*-value (1.6E-07) was smaller or equal to the original *P*-value (1.1E-06), yielding the permutation-derived *P*-value 0.001. For Iron-cT1, there were four such *P*-values, resulting in a permutation-derived *P*-value of 0.004 **(B)** The results after feature selection using different thresholds on correlation among the features. For each threshold *thr*, a minimal number of features were removed such that no pair of features remained with a correlation coefficient of a greater magnitude than threshold (Tegtmeyer, Arora et al., 2024).

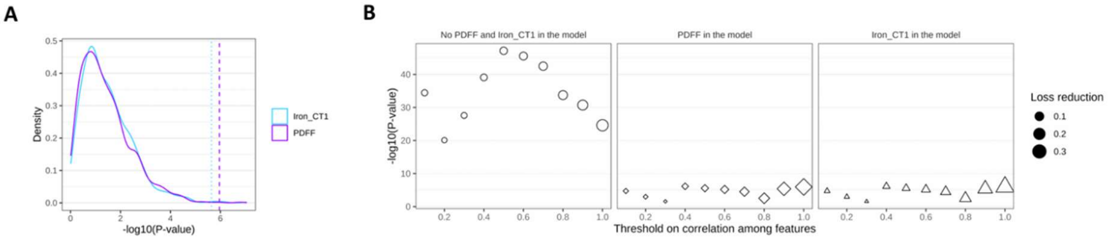

**Fig. S7. Manhattan plot of ancestor diseases with LR test p-value < 0.05.** Disease names are capped by 50 characters. The number of cases is shown in the brackets.

The tabular result is available at https://github.com/srm2022/iDWAS.

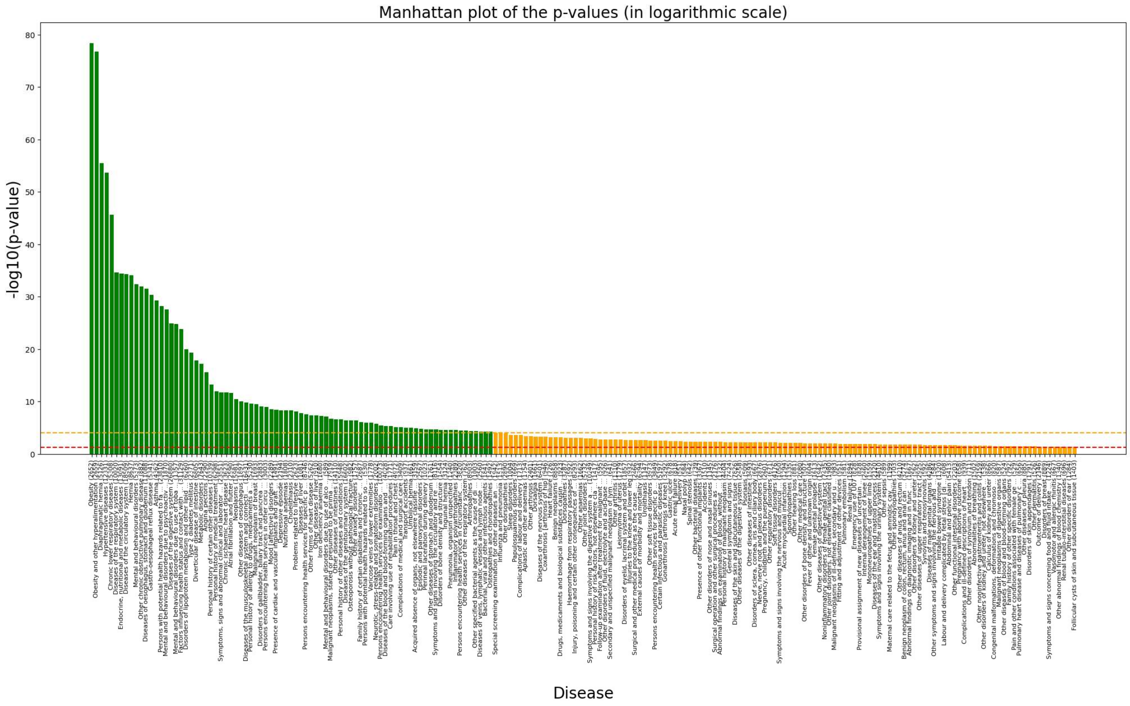


**Fig. S8. Manhattan plot of additional causes of death with LR test p-value < 0.05.** Disease names are capped by 50 characters. The number of cases is shown in the brackets. The tabular result is available at https://github.com/srm2022/iDWAS.

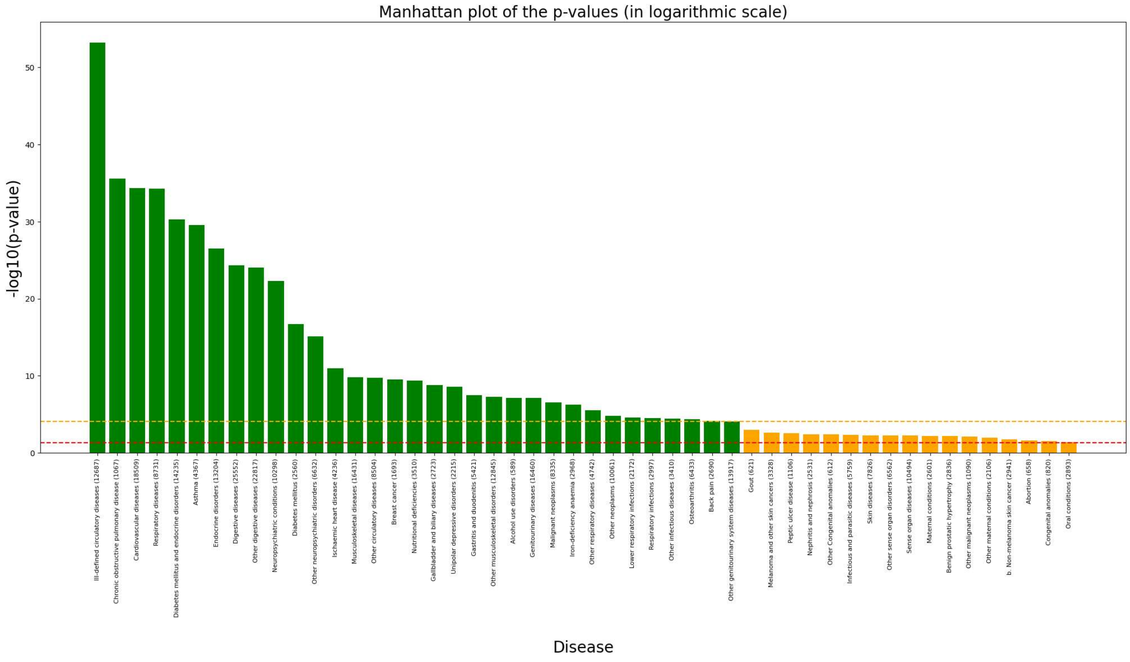

**Fig S9.** Diseases (N=77) with significant associations with the difference (Δp) in fitted probabilities between the proxy-MASH models with and without 512 MRI-derived features (using Bonferroni-corrected P-value threshold ($P$ < 0.05 / 562)).

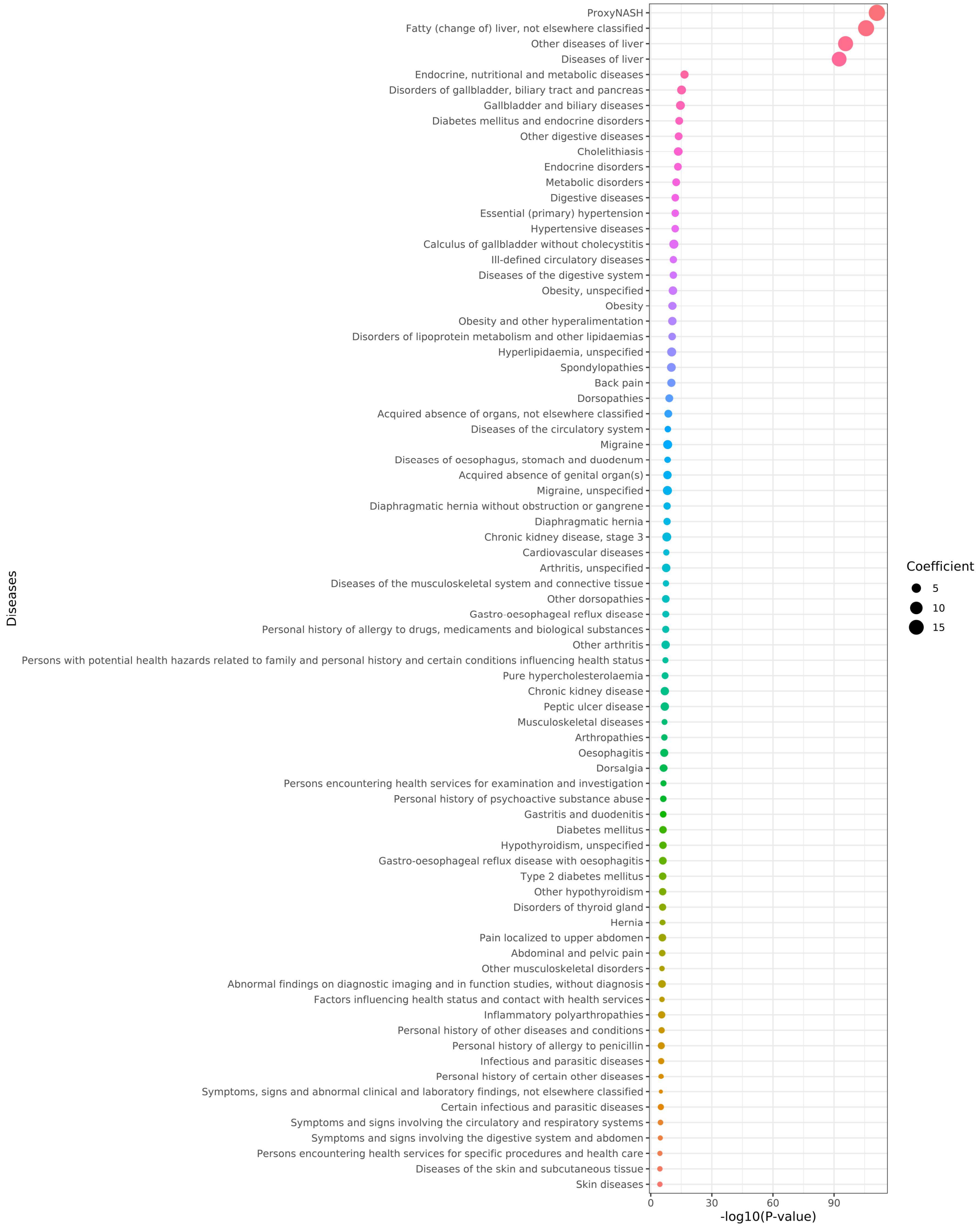
